# The architecture of allele-specific regulatory variant effects across five human genomes

**DOI:** 10.1101/2025.11.22.689315

**Authors:** Katherine Dura, Keith Siklenka, Kari Strouse, Shauna Morrow, William Majoros, Timothy E Reddy

## Abstract

The overwhelming majority of genetic associations with complex traits and disease involve non-coding genetic variants. Those variants are substantially enriched in gene regulatory elements, indicating that allelic impacts on gene regulatory element activity is a major contributor to those associations. As a step towards fine mapping the mechanisms within those associations, several studies have identified variants associated with gene expression and chromatin accessibility within those phenotypic associations. However, connecting those associations with functional impacts on gene regulatory element activity remains a major challenge. Here, we functionally measured allele-specific regulatory element activity across five human genomes using the genome-wide reporter assay STARR-seq. We identified tens of thousands of gene regulatory elements and estimated allele effects at ∼200,000 genetic variants therein, including ∼10,000 indels in our study population. Allelic effects on regulatory element activity correspond closely with predicted impacts on transcription factor binding motifs. The measured variant effects also allow us to fine map potential causal variants within eQTLs and chromatin QTLs from the same population. Together, these results provide an initial atlas of genome-wide variant effects across the human genome and demonstrate the potential for such approaches to prioritize causal variants for future mechanistic investigation.

## INTRODUCTION

Noncoding genetic variants play a major role in human health and disease. A common hypothesis is that many of those variants contribute to disease risk via their impacts on gene regulatory element activity. However, experimentally measuring those effects remains a longstanding challenge in human genetics^1^. Progress towards systematically measuring those non-coding variant effects has enormous potential to benefit human health. For example, targeted experimental studies of specific non-coding associations have highlighted novel genes and downstream pathways that have the potential to benefit diagnostics and therapeutics^2–4^.

A major step towards making the identification of noncoding disease mechanisms more routine is the development of high throughput approaches to systematically and comprehensively measure the effects of noncoding genetic variation on gene regulatory element activity. High throughput reporter assays such as STARR-seq and massively parallel reporter assays (MPRA) are promising technologies to achieve that goal. Those approaches have been used to measure genetic variant effects across individual gene regulatory elements^5,6^, across disease-associated genomic regions^4,7^, and across a single genome^8–11^. However, to the best of our knowledge, no studies have reported genome-wide measurements of the effects of genetic variants on gene regulatory element activity across multiple individuals.

As a step towards that goal, here we report the genome-wide effects of noncoding genetic variation on gene regulatory element activity across five individuals. To experimentally measure allele-specific regulatory element activity, we used a pooled STARR-seq approach in which we measure regulatory activity using fragmented genomic DNA from several individuals at once. Then, to estimate the effects of those variants on regulatory element activity, we used a recently developed Bayesian inference method^12^. Together, the study population contains millions of genetic variants. We identified tens of thousands of variants, including personal variants and insertions/deletions, within gene regulatory elements that had significant effects on regulatory element activity. Here, we present that catalog of regulatory variants. We assess the relative effects of different types of genetic variants, and attribute variant effects to specific transcription factor binding sequences. Finally, we demonstrate that the gene regulatory variants we identify can be used to fine map the effects of non-segregating genetic variants underlying genetic associations with traits and disease.

## RESULTS

### Genome-wide pooled STARR-seq to measure allele-specific gene regulatory element activity across five individuals

To measure the effects of genetic variants across the human genome on regulatory activity, we used the high-throughput reporter assay STARR-seq^13,14^. In STARR-seq assays, genomic DNA fragments are ligated into the 3’ untranslated regions of a synthetic reporter gene. From that location, the DNA fragments modulate activity of the upstream promoter. In doing so, the DNA fragments regulate their own rate of transcription into RNA. Regulatory element activity of each DNA fragment can thus be estimated by comparing the abundance of the fragment in transcribed RNA to the abundance of the fragment in the assay library. STARR-seq can also be used to assess allele-specific regulatory element activity by comparing the relative abundance of each allele in the RNA and assay library.

We measured regulatory element activity across the genomes of five Yoruba individuals from the 1000 Genome Project (1KGP) cohort (Fig. 1a). Together, their genomes include ∼14 million genetic variants that cover 80% of all common human genetic variants (minor allele frequency [MAF] > 5%), 26% of uncommon genetic variants (5% < MAF < 1%), and a substantial number of more rare and personal genetic variants (Fig. 1b). We assayed those variants using a pooled genome-wide STARR-seq approach, with five individual STARR-seq libraries combined together into a single assay. The average size of DNA fragments in the pooled library was 450 bp (Fig. 1c). That size is comparable to the average size of an open chromatin site. The pool includes ∼2 billion unique DNA fragments, and covers the human genome at ∼300X (Fig. 1d).

**Figure 1.**
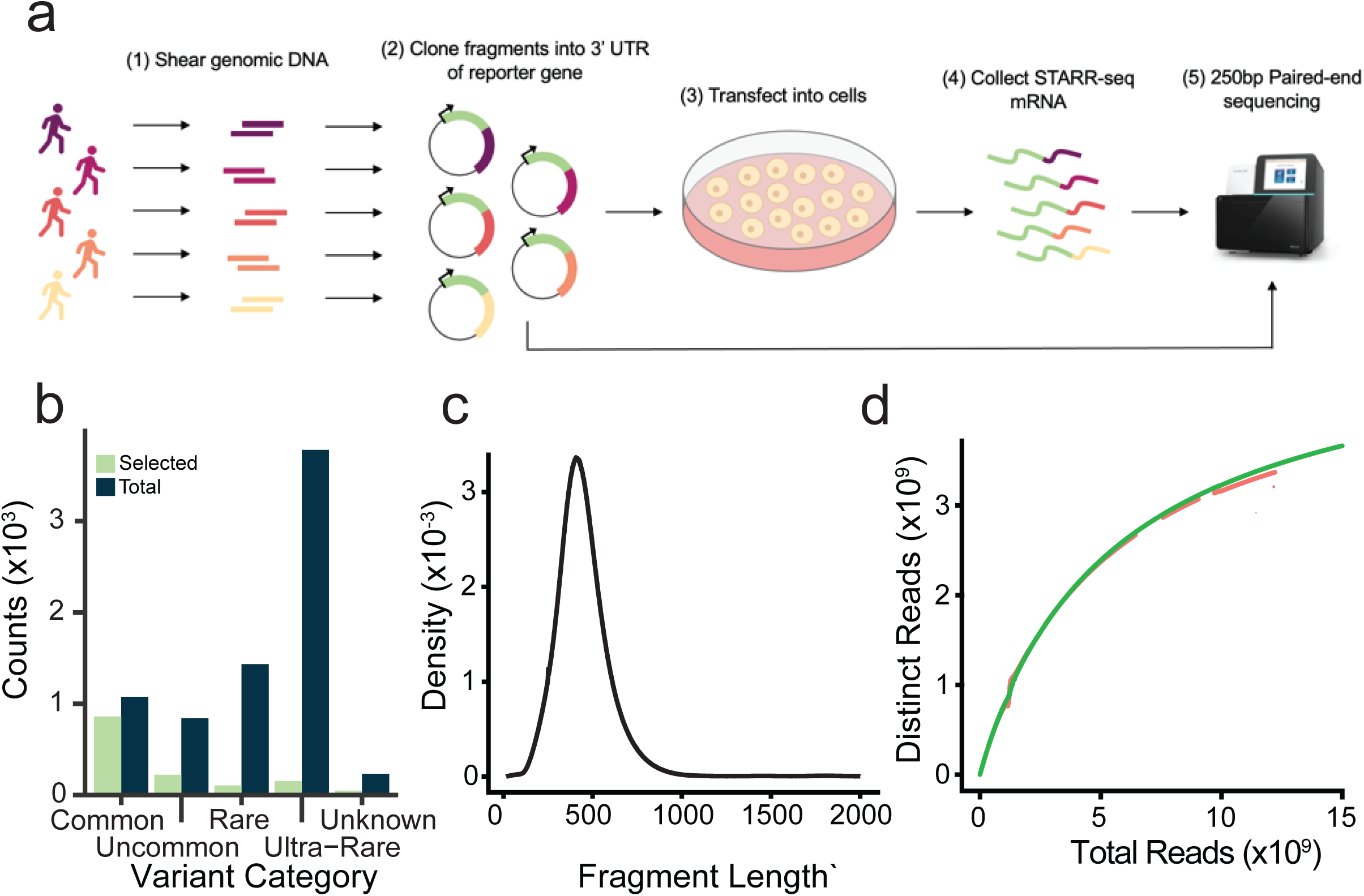
Experimental overview. A. Experimental design for the 5-sample pooled STARR-seq assay. Genomic DNA is sheared, and fragments are cloned into the 3’UTR of a reporter gene. The resulting library is transfected into K562 cells, and mRNA is collected from the cells and sequenced. B. Distribution of variants present in Thousand Genomes Project samples, for all samples (blue) and for samples included in this study (green) based on population frequency categories (Common: MAF>0.05. Uncommon: 0.01<MAF<0.05. Rare: 0.001<MAF<0.01. Ultra-Rare: MAF<0.001). C. Input library fragment length distribution. X-axis represents lengths of DNA fragments, and y-axis represents the density of the number of times a fragment exists at that length in the input DNA library. D. A per-replicate representation of extrapolated number of distinct reads in the experiments. X-axis represents total number of reads in the input DNA library, and the y-axis represents the expected number of distinct reads in the input DNA library.

### Genome-wide distribution of gene regulatory element activity

To estimate regulatory activity, we transfected the pooled assay library into K562 cells, and sequenced the cDNA from transcribed STARR-seq RNA. We completed the transfections in triplicate, each time transfecting ∼150 million K562 cells and sequencing to a depth of ∼150 million sequencing reads. Overall, there was strong correlation between the replicates and with the input library (Supplementary Fig. 1). The cDNA fragment size was reduced by ∼50 bp from the input library, likely due to the additional cycles of PCR preferentially amplifying shorter DNA fragments.

We used two complementary approaches to identify non-coding variants that impact regulatory activity. Because STARR-seq can also detect latent regulatory elements that are outside of open chromatin, we called regulatory element activity *de novo* from the STARR-seq data. We identified 56,570 genomic regions with regulatory activity at a 5% irreproducible discovery rate (IDR) (Supplementary Table 1). Of those, 56,340 (99.5%) activate gene expression and 230 (0.5%) repress gene expression.

Second, we hypothesized that regulatory variants will be enriched in open chromatin regions. We therefore estimated regulatory activity in open chromatin sites that serve as a baseline map of potential gene regulatory elements in the human genome. We focused our analysis on a leniently called set of 170,842 open chromatin sites identified with ATAC-seq in K562 cells. Those sites together cover ∼3.5% of the human genome. The *de novo* called regions were substantially enriched for open chromatin sites. For example, 18,436 (34%) of the de-novo called sites overlapped the 170,842 open chromatin sites we considered, a ∼10-fold enrichment over background. Across those sites, we found evidence of regulatory activity at 39,035 (23%) at a false discovery rate (FDR) < 5%. Most of those identified regulatory elements activate gene expression (N = 26,306, or 67%), while the remaining 12,729 (33%) repress gene expression (Fig. 2a; Supplementary Table 2).

**Figure 2.**
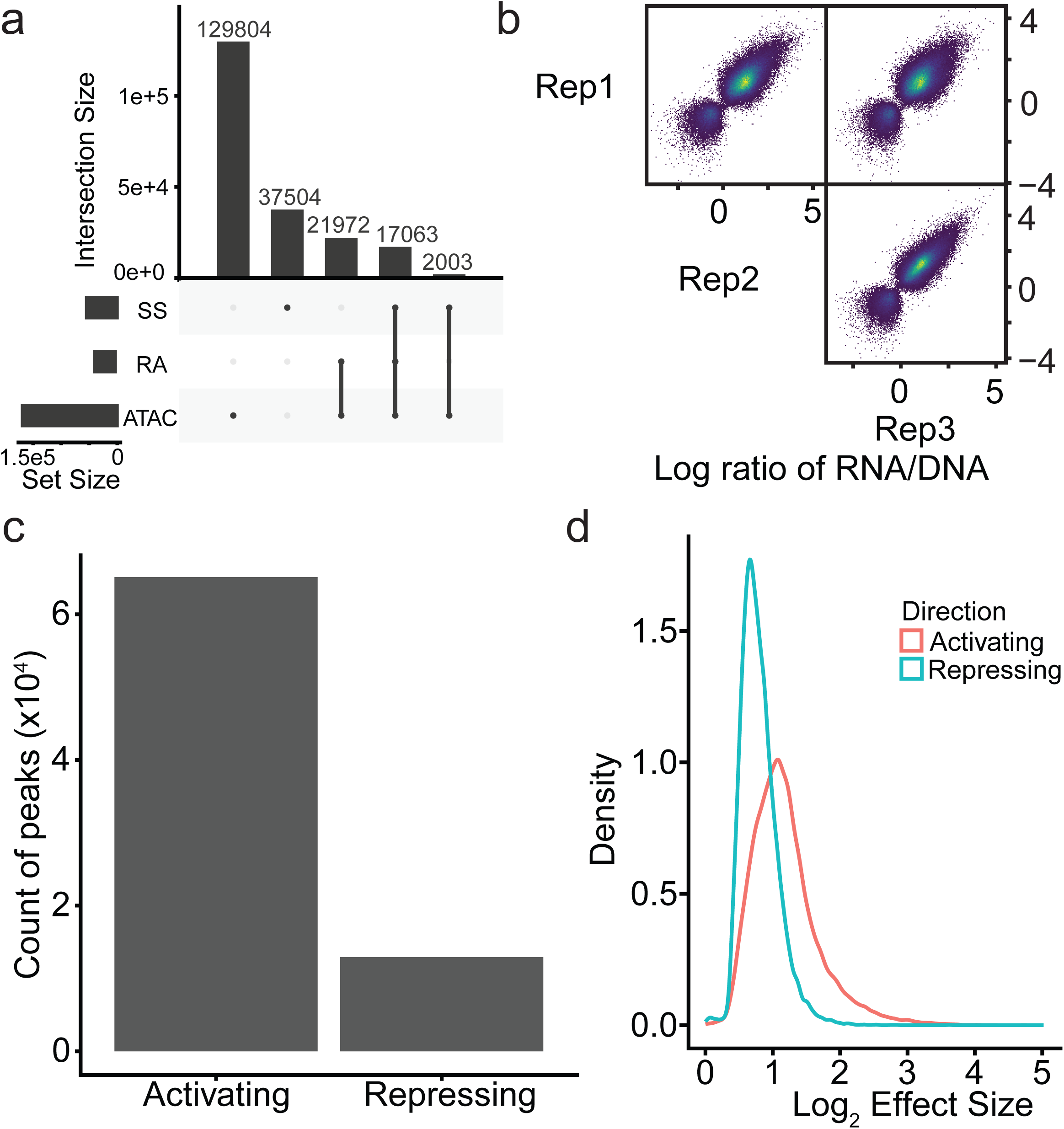
Properties of identified peak sets. A. Relative sizes of peak sets. The two main peak sets are ATAC-seq peaks (ATAC), some of which have regulatory activity (RA) in these experiments, and STARR-seq peaks (SS). Set size and intersection size indicate the number of peaks. Upset plot indicates the combinations of ATAC-seq peaks, ATAC-seq peaks with regulatory activity, and STARR-seq peaks in this dataset. B. Correlations of replicates in identified peak sets. The x- and y-axes are the log-transformed ratio of the RNA coverage to DNA coverage of the genomic regions in peaks. Each point represents a different peak in the genome. C. Proportions of activating and repressing peaks. The y-axis represents the count of peaks in each category. This dataset included 65,095 activating peaks, and 12,925 repressing peaks. D. Distribution of the relative magnitudes of effect sizes in activating and repressing peaks. The x-axis represents the absolute value of the log-transformed effect size. The y-axis represents the density of this effect size in the data.

Together, those two analyses give us a union set of 78,023 genomic regions with regulatory activity (Supplementary Table 3). We focus our downstream analysis on that set. Over that union set, estimated regulatory element activity was highly correlated between replicates (0.81 ≤ r ≤ 0.88. Fig. 2b); activating gene regulatory elements were more numerous than repressing gene regulatory elements (65,095 vs 12,925) (Fig. 2c); and activating gene regulatory elements had stronger effects than repressing elements (median effect size of 1.11 vs -0.75) (Fig. 2d).

### Genomic state at sites with gene regulatory activity

The identified gene regulatory elements were strongly supported by covalent histone modifications. Across the 65,095 genomic regions that activate gene expression, there was a strong central trend for covalent histone modifications canonically associated with gene activation, namely H3K27ac, H3K4me3, and H3K4me1 (Fig. 3a). H3K4me3 is considered to be more indicative of promoters, whereas H3K4me1 is indicative of enhancers^15^. That both marks were enriched in our results indicates that both enhancers and promoters have activity in our STARR-seq assays. In contrast, we did not observe strong enrichment for covalent histone modifications at repressive regions. Together, these results indicate that the activating and repressive sequences in STARR-seq have distinct chromatin states in the human genome, and that sequences with enhancer activity in STARR-seq also have evidence of active gene regulatory activity in the human genome.

**Figure 3.**
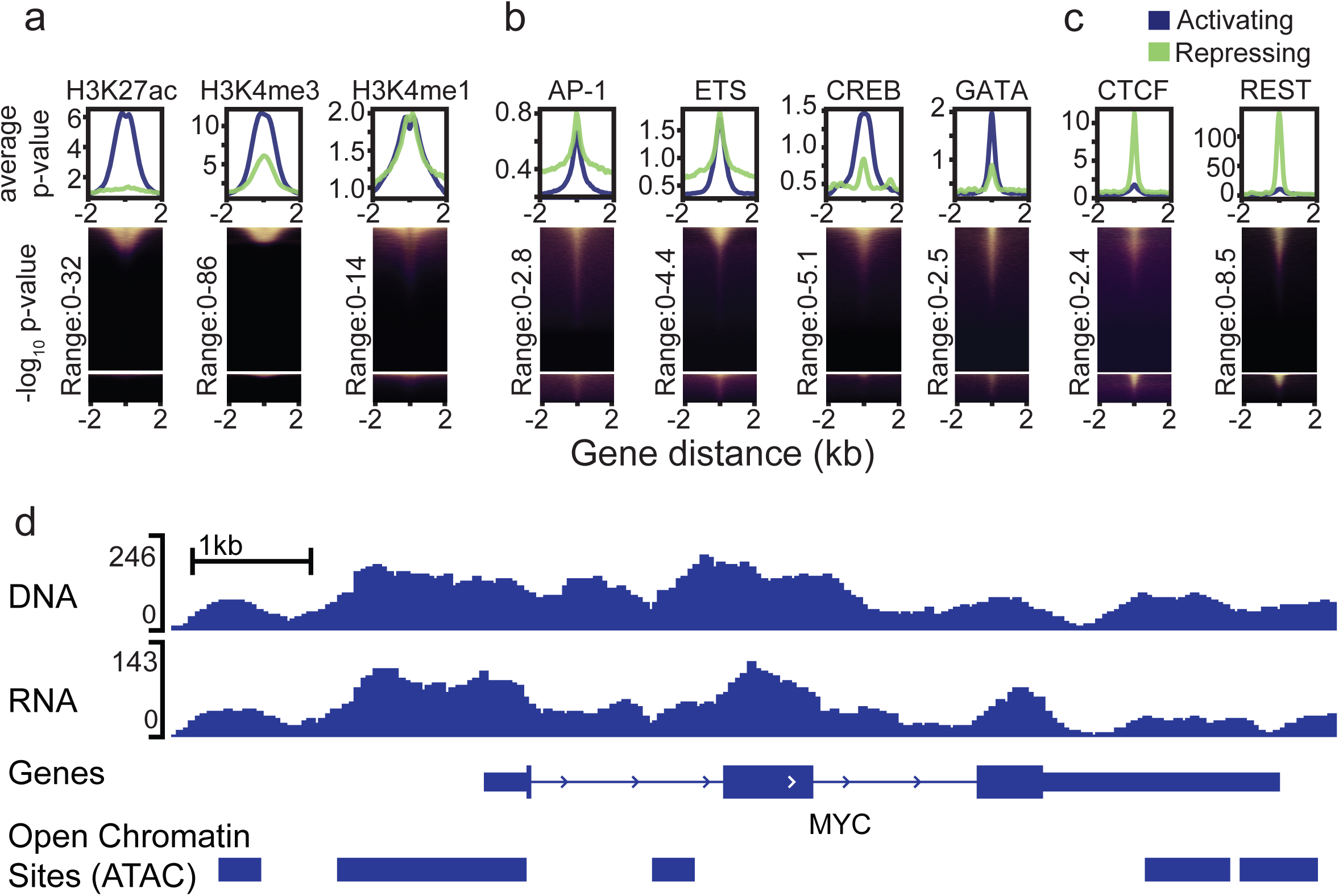
Identified regulatory marks within peaks. A. Histone modifications identified in peaks. The specific histone mark is listed. Heatmaps represent the relative -log10 p-values of the indicated histone mark occurring in each peak. Density plots represent the mean score across all activating peaks (blue) and repressing peaks (green) in the dataset. B. Transcription factor binding sites identified in peaks. The specific transcription factor is listed. Heatmaps represent the relative -log10 p-values of the indicated transcription factor occurring in each peak. Density plots represent the mean score across all activating peaks (blue) and repressing peaks (green) in the dataset. C. Repressing marks identified in peaks. The specific repressing mark is listed. Heatmaps represent the relative -log10 p-values of the indicated repressing mark occurring in each peak. Density plots represent the mean score across all activating peaks (blue) and repressing peaks (green) in the dataset. D. Browser track example of MYC locus. The coverage of all three DNA and RNA replicates are represented individu-ally. Beneath coverage tracks, the location of the MYC gene is represented. Below the gene representation, is a series of boxes representing each peak in the displayed region.

The activating and repressing activity identified via STARR-seq was also strongly supported by binding of activating and repressive transcription factors, respectively. Specifically, activating sites were commonly bound by activating transcription factors in K562 cells such as the AP1, ETS, CREB, and GATA families of transcription factors (Fig. 3b). In contrast, the 12,925 regions that repressed gene expression were distinctly enriched for binding by CTCF and REST, factors associated with repression and genome structure, respectively (Fig. 3c).

As further support that the identified regulatory effects are broadly consistent with gene regulatory activity in the genome, we compared our results to classes of candidate cis-regulatory elements (cCREs) defined by ENCODE. Specifically, we compared STARR-seq activity to version 4 of the cCRE catalog^16^. Of all the genomic regions that we identified as activating gene expression, 62% were also annotated as a proximal or distal enhancer-like region, resulting in a 4-fold enrichment over the background (Supplementary Table 4). Conversely, of the genomic regions that repress regulatory activity, 74% overlapped a CTCF-bound region, resulting in a 10-fold enrichment over the background (Supplementary Table 5). An example of the correspondence between STARR-seq activity and chromatin state in the MYC locus is shown in Figure 3d.

### Estimation of genetic impacts on regulatory element activity

We next estimated allele specific regulatory activity across our cohort. There are 14 million genetic variants in our study population. Those variants were observed at their expected frequencies in the pooled STARR-seq input library, indicating a lack of allelic bias during library construction (Fig. 4a). Importantly, because we pooled genomes of five individuals in our assay library, the minimum allele frequency is 10%, even for personal and ultra-rare variants in the human population. Because of that floor on the minimum allele frequency, we observed both alleles of nearly all variants in called regulatory elements at high frequencies. For example, in the set of STARR-seq regulatory elements described above, there were on average 50 reads for alternate alleles that were present in our final library at 10% (Fig. 4b).

**Figure 4.**
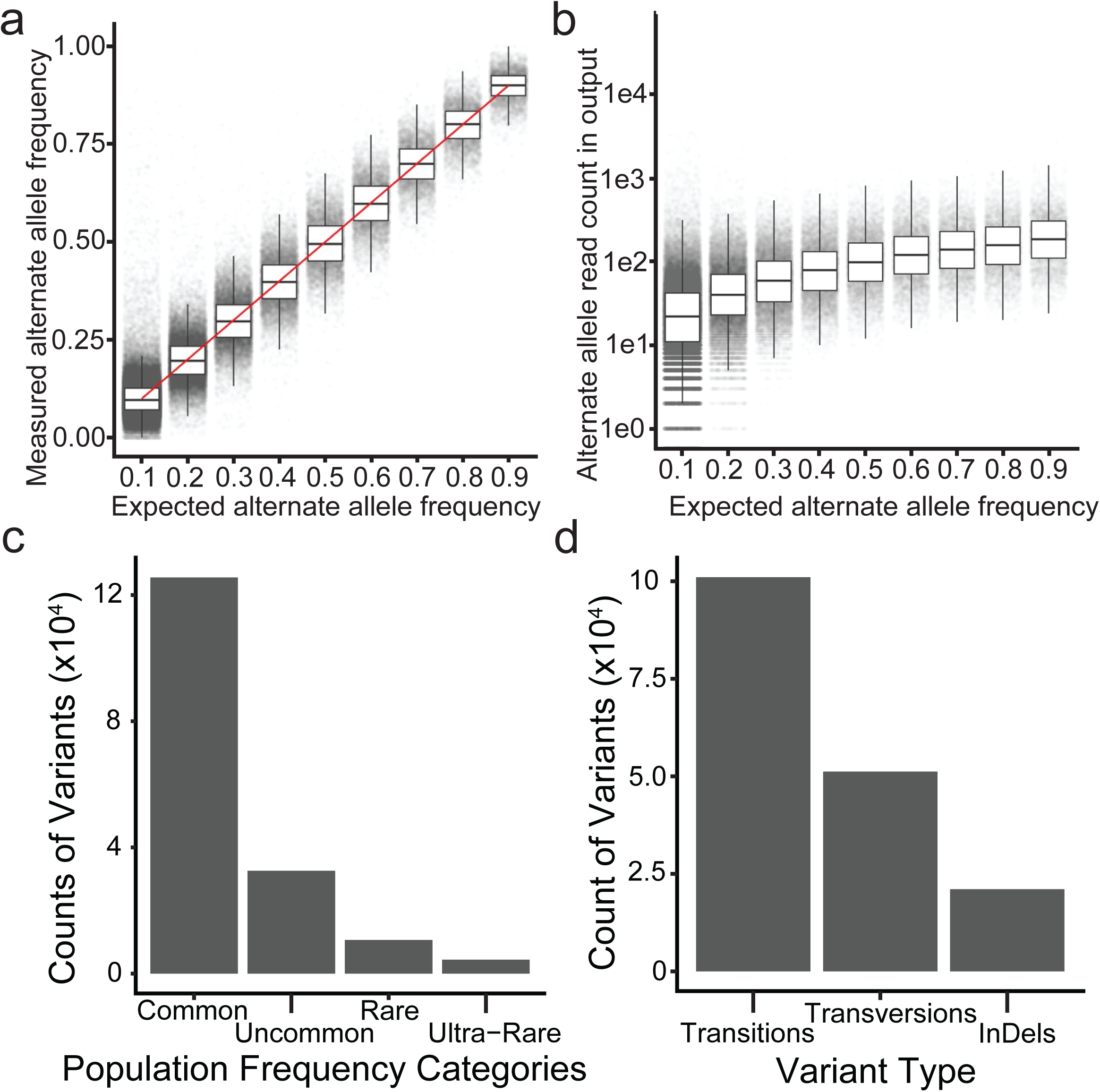
Properties of individual variants A. Measured alternate allele frequencies of assayed variants. Each point represents a different assayed variant. The x-axis represents the expected alternate allele frequency in the pool. The y-axis represents the measured alternate allele frequency in the DNA library. The red line is the y=x line. B. Measured RNA coverage of assayed variants. Each point represents a different assayed variant. The x-axis represents the expected alternate allele frequency in the pool. The y-axis represents the coverage of the alternate allele in the RNA library. C. Proportions of variants in different population frequency categories. The y-axis represents the counts of variant population allele frequency categories in this dataset. This dataset includes 126,249 common variants, 32,774 uncommon variants, 10,731 rare variants, and 4,474 ultra-rare variants. D. Proportions of different categories of variant types. The y-axis represents the counts of different variant types in this dataset. This dataset includes 101,472 transitions, 51,438 transversions, and 21,318 indels.

We estimated allelic biases in regulatory element activity for all of those variants using a Bayesian hierarchical model designed for that purpose (Supplementary Table 6)^12^. Overall, we tested 187,493 variants in called regulatory regions that had more than 5 reads in the sequencing of our input library and in each replicate output library. We found 5,393 significant allelic effects on regulatory element activity. Allelic effects were typically small and had low posterior probabilities of having an effect, indicating that most genetic variants do not impact regulatory element activity in our assays (Table 1).

**Table 1.**

| Read Count | Total Variants | $P_{reg} > 0.8$ count | $P_{reg} > 0.8$ percentage | $P_{reg} > 0.9$ count | $P_{reg} > 0.9$ percentage |
| --- | --- | --- | --- | --- | --- |
| 5 | 187,493 | 10,315 | 5.502% | 5,393 | 2.876% |
| 10 | 178,651 | 9,852 | 5.515% | 5,167 | 2.892% |
| 25 | 146,543 | 7,916 | 5.402 | 4,206 | 2.870% |
| 50 | 96,720 | 4,890 | 5.065% | 2,590 | 2.678% |

Most of the high confidence variant effects were found with common alleles (72%), followed by uncommon alleles (19%), rare alleles (6%), and ultra-rare alleles (3%) when using GnomAD to estimate the population frequency of alleles in individuals with African ancestry (Fig. 4c). The detected variants include transitions (58%), transversions (30%) and indels (12%) (Fig. 4d).

### Impact of rare variants on regulatory activity

Several studies have suggested that rare and ultra-rare non-coding genetic variants have greater impacts on regulatory element activity, potentially reflecting either selection against such large variant effects or new variants in the human population that have yet to be selected against. A rolling average of effect sizes across variants ordered by their minor allele frequency in the African population also demonstrates a trend where more rare variants have greater effects on regulatory element activity (Fig. 5a). However, that increased effect size was explained by low minor allele counts in the sample. Specifically, there was no significant association between allele frequency and the size of the effect on regulatory element activity when comparing alleles with a minor allele count of one in this study independent of the population allele frequency (p=0.309) (Fig. 5b).

**Figure 5.**
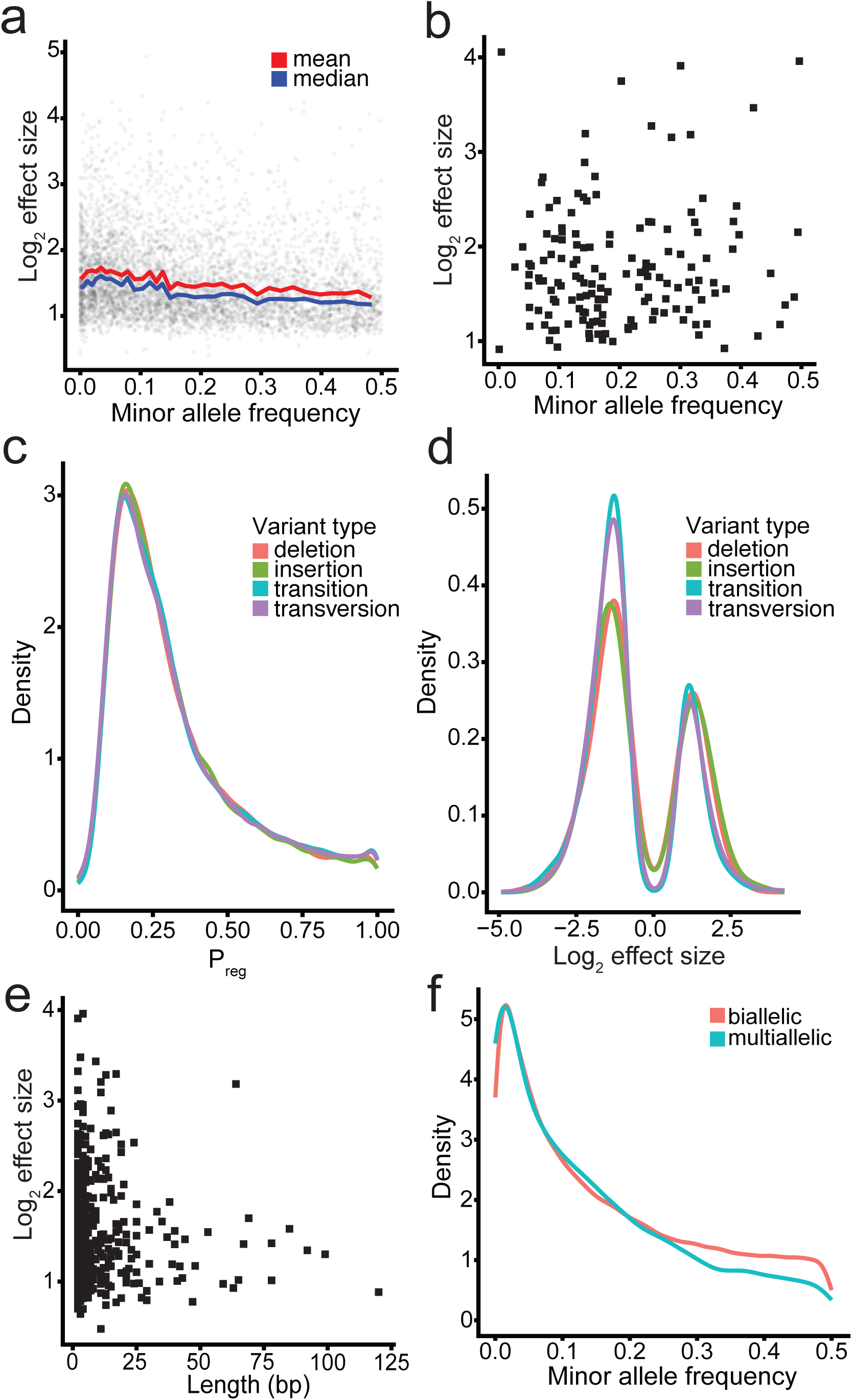
Properties of variant effect sizes A. Comparison of variant minor allele frequency and variant effect size for all assayed variants. Each point represents a different assayed variant. The x-axis represents the minor allele frequency of the variant in the population. The y-axis represents the log-transformed STARR-seq effect size. The rolling mean is represented in red, and the rolling median of this data is represented in blue. B. Comparison of variant minor allele frequency and variant effect size when the study frequency is 0.1. Each point represents a different variant that occurs in the dataset with an expected frequency of 0.1. The x-axis represents the minor allele frequency of the variant in the population. The y-axis represents the log-transformed STARR-seq effect size. C. Comparison of posterior probability of variant effects among different variant types. Deletions are represented in pink; insertions are represented in green; transitions are represented in blue; and transversions are represented in purple. The x-axis represents the posterior probability that a variant has an effect given the results of the STARR-seq assay. The y-axis represents the density of this posterior probability in the data. D. Comparison of variant effects among different variant types. Deletions are represented in pink; insertions are represented in green; transitions are represented in blue; and transversions are represented in purple. The x-axis represents the log-transformed STARR-seq effect size. The y-axis represents the density of this effect size in the data. E. Comparison of indel length and variant effect size. Each point represents a different variant. The x-axis represents the length of the insertion or deletion in bases. The y-axis represents the absolute value of the log-transformed STARR-seq effect size. F. Comparison of allele frequency distribution of biallelic variants and multiallelic variants. The biallelic distribution is represented in pink, and the multiallelic distribution is represented in blue. X-axis represents the minor allele frequency of the variants. The y-axis represents the density of this minor allele frequency in the data.

### Effects of insertions and deletions on regulatory element activity

Effects of insertions and deletions (indels) have been a historically challenging to assay, in part due to the difficulty in correctly aligning to indels and identifying them. We hypothesized indels will have greater effects because they can more substantially impact the structure of transcription factor binding sequences than single nucleotide substitutions. In addition, larger indels may add or remove entire binding sites. To test that hypothesis, we evaluate the effects of indels on regulatory element activity genome-wide. Due to challenges aligning short sequencing reads to insertions and deletions, we developed a custom sequence alignment method to improve detection of sequences containing insertions and deletions in our study samples. Briefly, we create local genome sequences that include all haplotypes of regions with genetic variation in our sample, and we then align to those haplotypes using strict alignment quality criteria. That approach is an elaboration of previous related methods used for allele-specific alignment to the genome of one individual^17,18^.

Across our study pool, we estimated the effects of 21,063 indels on regulatory element activity in regulatory regions. Indels were as likely as single nucleotide variants to have a significant effect on regulatory element activity (Fig. 5c). Specifically, 2.7% of tested indels (N = 575) had significant effects on regulatory element activity, compared to 3% for SNVs. Overall, when indels had a significant effect on regulatory element activity, the magnitude of the effects was comparable to that of SNVs; however, indels were slightly less likely to have negative effects on regulatory element activity (Fig. 5d). Short (<5 bp) indels had a similar distribution of effect sizes, although the small number of large indels assayed makes that comparison challenging (Fig. 5e). Nevertheless, we found that indels commonly and substantially disrupt regulatory element activity across the human population. Those results agree with a previous association study showing indels have similar correlations to SNPs with a range of anthropomorphic and blood cell traits^19^. That SNVs and indels have similar effect magnitudes suggests that the effects of noncoding variants depend less on the type of variant and more on the specific DNA sequence changes affected.

### Investigation of multiallelic sites in regulatory elements

2.24% of variable sites in the human genome are multiallelic, most of which are variable length insertions and deletions^20^. With multiple samples used in this experiment, we were able to examine the effects of genetic variants that have multiple alternate alleles. Specifically, we estimated the effect of multiallelic sites relative to all other alleles using BIRD. There was a total of 2,576 multiallelic sites, with 5,868 non-reference alleles in the peak set described above, with up to 6 non-reference alleles at a single site (compared to 167,497 of bi-allelic sites). As expected, the population frequencies of these multiallelic variants were lower than those of biallelic sites (Fig. 5e). When comparing the significance of variants across these sites, we found significant effects at 101 sites (4%). In all but five cases, we detect a significant effect for only one non-reference allele. In the five cases where we detect effects for multiple non-reference alleles, the variants have two alternate alleles and both are significant. In the cases where there is a variable length repetitive element, increasing the length of the repetitive element resulted in a lower transcription rate. In cases where there is a SNV, the alternate allele results in a lower rate of transcription according to the STARR-seq effect size.

### Observation of Transcription Factor Effects with STARR-seq Variants

Based on the above results, we next sought to identify specific transcription factor binding sequences that were disrupted by non-coding genetic variants. To do so, we estimate how those variants impact predicted transcription factor binding using motifbreakR^21^. We identified 329,146 instances of transcription factor binding sequences from the 393 motifs in the HOCOMOCO database. Clustering together overlapping instances of highly similar motifs^22^, we arrived at a final test set of 207,556 instances of 168 motif clusters.

To identify variant effects at specific positions of specific motifs, we developed a correlation-based approach. In that approach, we compared estimated allele effect sizes from motifbreakR to the allele effect size from the STARR-seq assay. We found four motif clusters had high concordance between the predicted TF binding and regulatory activity: AP-1, GATA, CREB, and ETS (Fig. 6a, Supplementary Fig. 2-5). Each motif cluster represents a family of up to dozens of individual transcription factors that bind similar genomic sequences. The AP-1 family is a broadly acting transcription factor that has roles in a wide range of cell types. Meanwhile, multiple GATA, CREB, and ETS factors have been identified as having key roles in hematopoietic cell differentiation. The identification of those motifs here corresponds to the fact that we measured variant effects in K562 cells, a leukemia cell line. We also estimated whether specific positions in the TF binding motif were enriched or depleted for significant STARR-seq variant effects. Overall, we found that positions in the motif with greater information content were more likely to have a significant variant effect, as expected (Fig. 6b, 6c, p<2^-16^). Together, these results indicate that the STARR-seq assays capture both variants that affect general transcription factor binding, and variants that affect cell-type specific gene regulatory mechanisms.

**Figure 6.**
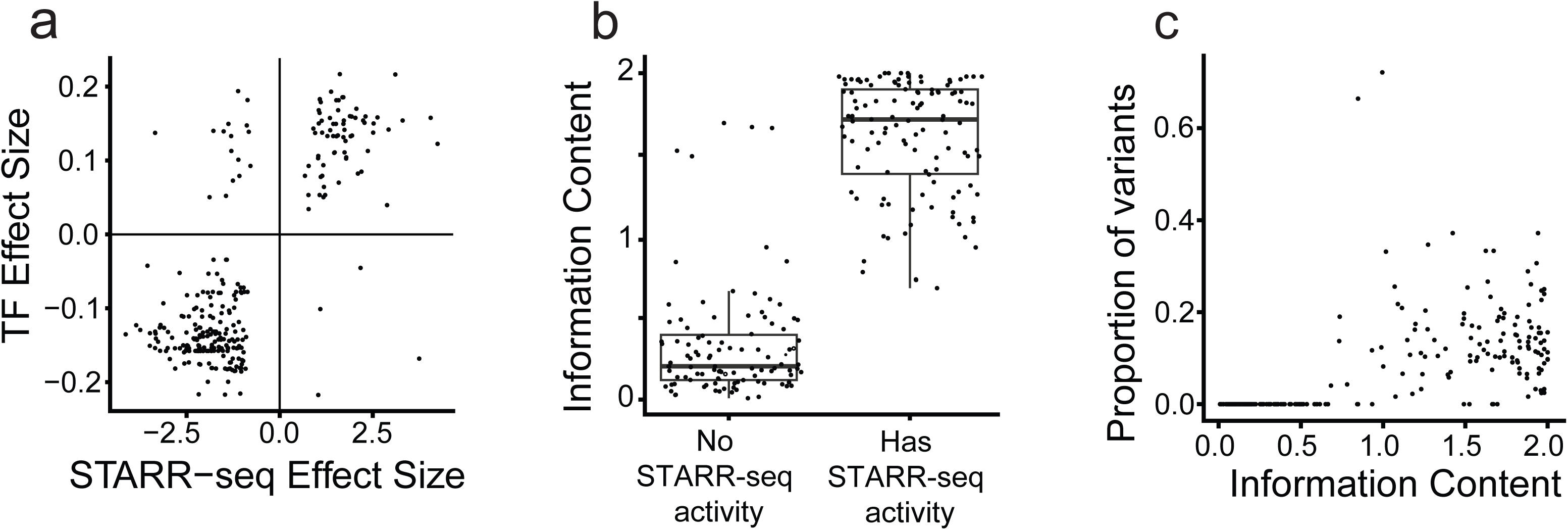
Comparisons to TF binding sites A. Correlation between STARR-seq effect size and predicted effect size caused by changes to transcription factor binding sites. Each point represents a different variant. X-axis represents the aggregated log-transformed effect size from STARR seq for all variants in the open chromatin region. Y-axis represents the predicted change in TF binding caused by that variant. B. Comparison of information content of positions in transcription factor binding sites that include STARR-seq variants, and those that do not include STARR-seq variants. Each point represents a different position in the assayed PWMs. PWM positions are divided between those with no STARR-seq variants, and those with at least one STARR-seq variant. C. Comparison between likelihood of variants at transcription factor binding site positions and the information content at those positions. Each point represents a different position in the assayed PWMs. X-axis represents the information content at that position. Y-axis represents the percentage of variants that occur in the correct position in that motif.

### Correlations with QTLs

A key step in resolving the genetic basis for human traits and diseases is fine mapping genetic associations to individual causal variants. To identify potential causal variants in this population-wide STARR-seq study, we compared our estimates of the effects of genetic variants on regulatory element activity to the effects from genetic association analyses. We focused on associations from the African Functional Genomics Resource (AFGR) because the study populations have matched ancestries and all five individuals studied here are included in that study. Briefly, the AFGR study investigated genetic associations with gene expression (i.e. eQTLs, N = 593 individuals) and chromatin accessibility (i.e. caQTLs, N = 100 individuals) across diverse African individuals^23^.

We detected significant correlation between the effects of genetic variants on regulatory element activity and the effects of caQTLs from the AFGR datasets (correlation coefficient is 0.675) (Fig. 7a). These correlations were found to increase as the posterior probability of the STARR-seq variant increased (Fig. 7b). We were not able to find significant correlations between eQTLs and STARR-seq variants, and the majority of variant pairs in similar regions had effect sizes that did not match in direction. We hypothesized that was because most eQTL associations are due to linkage disequilibrium, and that only a small fraction of regulatory variants have a functional effect. In both cases, there were fewer functional variants in the STARR-seq dataset for a single open chromatin region than QTLs. When considering all variants regardless of the strength of the allelic effects, each caQTL and each eQTL could have over 100 associated STARR-seq variants. After taking into account only the variants that were most likely to have regulatory activity, each eQTL had at most two associated STARR-seq variants, and each caQTL had at most four associated variants, greatly reducing the potential number of impactful variants in these regions and indicating a potential of fine-mapping of regulatory regions with this data type.

**Figure 7.**
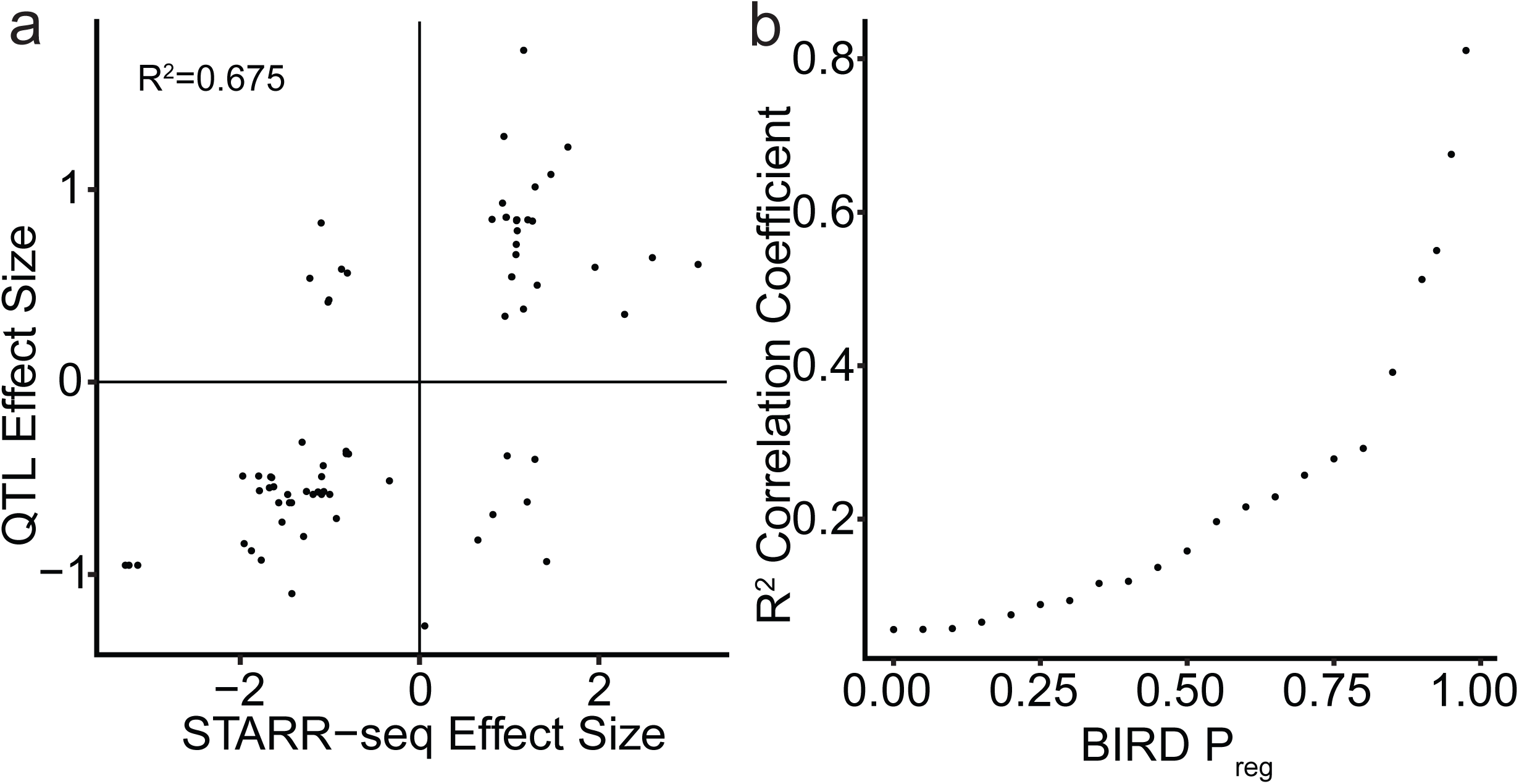
Comparisons between STARR-seq variants and QTLs. A. Correlation between QTL effect size and aggregated STARR-seq effect size across the corresponding open chromatin region. X-axis represents the aggregated log-transformed effect size from STARR seq for all variants in the open chromatin region. Y-axis represents the log-transformed effect size for the associated QTL. B. Demonstration of increasing correlation with QTLs when confidence in STARR-seq variant effects increases. X-axis represents the minimum posterior probability that the STARR-seq variants have an effect. Y-axis represents the R2 correlation coefficient between QTL effect sizes and aggregated STARR-seq effect sizes with the correspond-ing thresholding.

## DISCUSSION

Genetic impacts on gene regulatory activity are thought to contribute to a wide range of traits and diseases. Here, we completed a genome-wide assessment of noncoding genetic variants effects on gene regulatory element activity across the genomes of five individuals. That enabled us to estimate the effects of hundreds of thousands of non-coding variants including rare variants and indels; to relate those variant effects to changes in transcription factor binding; and to fine map within chromatin QTLs to prioritize subsets of highly functional variants. This work builds on previous studies that have experimentally fine mapped a single trait or disease, typically focusing on a few strongly associated. Larger scale studies have used MPRA to identify common genetic variants that modulate expression. Those studies also found enrichment for putative causal variants within those sets. Here, we present a general genome-wide approach that includes more types of genetic variants and longer assay fragments.

One of the key challenges is aligning long reads containing many variants in a way that does not create a reference bias^24^. Previous studies have used a range of approaches including filtering reads^25^ and creating local diploid sequences for alignment^17,18^. Since those approaches were published, the read length from high-throughput sequencing has grown from ∼75 bp reads to the 250 paired end reads we used here. Here, we adapted the diploid sequence approach to accommodate a pool of five individuals and longer sequencing reads. To do so, we aligned against several haplotypes at once. However, to avoid combinatorial explosion of potential haplotypes, we also needed to limit the length and the number of variants in locally variable regions. Ultimately, that alignment method was particularly important to accurately estimate the effects of both rare variants and indels, which have previously been understudied.

The ability to systematically investigate the effects of rare genetic variants here allowed us to reassess whether rare variants are more or less likely to impact gene regulatory element activity. Some studies have suggested that rare variants are more likely to have functional effects because they are actively selected against^26^. Our study design and analysis approaches enabled us to assay 15,205 rare and personal variants. While uncommon and rare variants had greater effects on regulatory activity on average, those effects were explained by the low frequency with which the corresponding alternate allele was sequenced. That result conflicts with a previous report that rare variants have greater effects^12^, however that study design did not allow disentangling effect sizes and allele counts. Taken together, we suggest here that rare variants generally do not have greater effects on gene regulatory element activity than common variants; or that if there is such an effect, the size of that effect is small on average.

We do however find that certain types of variants do have greater effects. In particular, short insertions and deletions and variants that disrupt high-information-content positions in transcription factor binding motifs had greater effects overall. Moreover, the effect of transcription factor motif disruption was particularly pronounced for a small number of factors known to be important in the K562 cell line^27,28^. One possibility is that those transcription factor binding motifs have overall greater information content or are especially highly expressed, making for larger effects that are also easier to detect. While much remains to be understood, it is possible that strict prioritization of variants within well-defined positions in highly expressed transcription factors may be an effective strategy to prioritize functional follow-up in future studies.

Finally, variant effects estimated via STARR-seq correlated with the QTLs of molecular traits, particularly when the QTL study assessed individuals with the same ancestry. That correlation was also particularly pronounced for chromatin QTLs which are typically evaluated more locally than eQTLs. Because there is functional information only present in the STARR-seq assay, we suggest that STARR-seq variants can therefore fine-map within those chromatin QTLs and can aid in the discovery of causal variants underlying molecular associations.

## METHODS

### Whole-genome STARR-seq input library generation

To develop our genome-wide and population-wide pooling approach to STARR-seq, five samples were selected from 1000 Genomes Project Yoruba population. To build STARR-seq input libraries, each genome was randomly sheared to 500 bp and those fragments were cloned into the 3’UTR of the STARR-seq reporter vector. An input library for each genome was built separately, and those five libraries were pooled to create the first pooled STARR-seq input library. Each library contains a few billion unique fragments, resulting in approximately 300X genome coverage.

The protocol used for STARR-seq input libraries is documented on the ENCODE portal^16^ and as follows. The original protocol can be followed here: https://www.encodeproject.org/documents/d0e78573-c6ca-4434-b736-3f628059bed9/@@download/attachment/wgSTARRseq_ENCODE_KPS_2021.pdf

The DNA fragments were sheared to 500 bp. The libraries were prepared with the NEBNext DNA Library Prep Master Mix Set for Illumina (Cat. #E6040L) through the End-Repair and A-tailing steps with few modifications. First, the ligation step was performed with a 15 μM STARR-seq adapter. After the first SPRI cleanup after the ligation step, the 500 bp insert SPRI size selection was performed. The STARR-seq library was amplified with the KAPA HiFi HotStart PCR Kit with the following reaction conditions: 1 μL HiFi polymerase, 10 μL GC 5x buffer, 10 μL 10 μM primer mix, 1.5 μL 10 μM dNTPs, 25 μL DNA library, 2.5 μL water. The PCR was performed with the following conditions: 3 minutes held at 95 °C, followed by 10 cycles of 20 seconds held at 98 °C, 15 seconds at 63 °C, and 1 minute at 72 °C. Following the cycles, the samples were kept at 72 °C for 5 minutes, then held at 4 °C. The enriched libraries were pooled, and a 0.9x SPRI cleanup was performed, and eluted in 100 μL. Samples were stored at 20 °C.

Next, these vectors were linearized. STARR-seq plasmid was digested with Agal and Sall. Products were run on a large 1 % agarose gel and a 1 kb ladder. The gel was stained with SybrSafe, and the large band was excised. The fragment was recovered with a GeneJet Gel Extraction Kit.

A Gibson Assembly was performed with 0.2 pmols DNA fragments, a 3:1 insert:vector molar ratio, 10 μL NEBuilder HiFi DNA Assembly MM, and up to 20 μL water. This reaction was incubated at 50 °C for 15-30 minutes, and immediately placed on ice. Sample reactions were pooled to begin ethanol precipitation by adding 0.1X volume 3 M NaOAc and 2.5X volume very cold 100 % EtOH. Samples were stored at -20 °C overnight.

Samples were centrifuged at 16,000xg for 30 minutes at 4 °C. Pellets were washed with 500 μL cold 70% EtOH, and centrifuged at 16,000xg for 10 min at 4 °C. This step was repeated once. DNA pellets air dried for a few minutes, then were resuspended in 40 μL water. Gibson products were split across 8 cuvettes per genome, each with 300 μL of in-house grown and frozen Endura electrocompetent cells. Samples were transformed by electroporation with the E. coli 2 mm 3 kV settings. 1 mL Lucigen Recovery media warmed to 37 °C was added. Cuvettes were washed 4X and collected in a tube for a total of 5 mL. Samples were shaken for 1 hour at 37 °C. Diluted culture was plated to determine transformation efficiency and estimate library diversity. Transformations were pooled into 1 L LB with carbenicillin and shaken for 8 hours at 37 °C. Transformed cells were pelleted at 5000xg for 15 minutes at 4 °C and stored at -20 °C.

DNA plasmid libraries were harvested using 1 Giga-prep column for 2 flasks. STARR-seq input sequencing libraries were amplified off of plasmid DNAs using the Kapa HiFi HotStart PCR Kit. The reaction conditions for each sample were as follows: 20 ng plasmid DNA, 5 μL GC buffer, 0.75 μL 10 mM dNTPs, 1.5 μL 10 μM primers, 0.5 μL polymerase, and water up to 25 μL. The cycling conditions for the PCR were as follows: 98 °C for 45 seconds, followed by 9 cycles of 15 seconds at 98 °C, 30 seconds at 64.5 °C, and 1 minute 10 seconds at 72 °C. Following the cycles, samples were kept at 72 °C for one minute, then held at 4 °C. Double-sided bead cleanup was performed (0.5X, 0.9X), then the size distribution was assessed on a Tape Station.

### Whole-genome STARR-seq output library generation

The STARR-seq library was transfected into K562 cells. Cells were harvested 6 hours post-transfection. RLT buffer was added to the cells with 1% beta-mercaptoethanol, and cells were vortexed. The cells were flash frozen in 50mL conical tubes, and stored at -80 °C.

Thawed samples were passed through an 18- to 20-gauge needle 10 times. RNAs were isolated through Qiagen RNeasy midi-columns, while performing the on-column DNase step. Samples were Eluted twice using 125 and 75 μL RNase-Free water. RNA concentration was quantified and R.I.N values were assessed. 1 μL RNase Block was added, and cells were stored at -80 °C

The volume of RNA was adjusted with 10 mM Tris-HCl buffer (pH 7.5) to 100 μL. RNA was heated to 65 °C for 2 minutes, then placed on ice.

200 μL (1.0 mg) of resuspended Dynabeads were transferred to a μ centrifuge tube. The tube was placed on a magnet, and the supernatant was discarded. The tube was removed from the magnet, and 100 μL 2x Binding Buffer was added (20 mM Tris-HCl Ph 7.5, 1.0M LiCl, 2 mM EDTA). The tube was returned to the magnet, and the supernatant was discarded. The tube was removed from the magnet again, and an additional 100 μL of 2x Binding Buffer was added to the Dynabeads, followed by the RNA. Samples were rotated at room temperature for 5 minutes. The tube was returned to the magnet, and the supernatant was discarded. The tube was removed from the magnet, and washed twice with 200 μL Washing Buffer B (10 mM Tris-HCl Ph 7.5, 0.15 M LiCl, 1 mM EDTA). The sample was eluted with 100 μL of 10 mM Tris-HCl, pH 7.5, and heated to 75 °C for 2 minutes, and the tube was immediately placed on the magnet. Eluted mRNA was transferred to a new RNase-free tube on ice.

Dynabeads were washed twice with 50 μL of 2x Binding Buffer and 50 μL RNAse-free water. 100 μL 2x Binding Buffer was added to the beads after washing.

Eluted RNA was heated to 65 °C for 2 minutes, then placed on ice. 100 μL of RNA was added to the Dynabeads suspension, and rotated for 5 minutes at room temperature. The tube was placed on the magnet, and the supernatant was discarded. The tube was removed from the magnet and washed with the mRNA-bead complex twice with 200 μL Washing Buffer B. Samples were eluted with 40 μL of 10 mM Tris-HCl, pH 7.5, and heated to 75 °C for 2 minutes before being placed immediately on the magnet. 40 μL quantities of mRNA were transferred to new RNase-free tubes on ice.

To each 40 μL mRNA sample, 10 μL DNase digestion mix was added (5 μL 10X TURBO DNase Buffer, 1 μL TURBO DNase, 1 μL RNAse Block, and 2 μL water), and incubated at 37 °C for 30 minutes. A 5 μL DNAse Inactivation Reagent was added, and the sample was incubated for 5 minutes at room temperature. Samples were centrifuged at 10,000 x g for 1.5 minutes. The 40 μL RNA was transferred to a fresh tube, making sure that the bead slurry did not carry over. We added 35 μL of water to each tube with beads to perform back extraction with a final volume of ∼50 μL. Samples were centrifuged again at 10,000 x g for 1.5 minutes, and 27 μL of supernatant were transferred to matched supernatant from the first spin, making sure the slurry did not carry over. 1 μL of Agilent RNAse block (40 units/μL) were added to the samples, then vortexed to mix for a final volume of 68 μL.

First-strand cDNA synthesis was performed using the SuperScript III system and reporter-specific primers. 10 μL of the first master mix was added to 68 μL of each sample (3 μL of 4 μM RT_UMI Primer, 6 μL of 10 mM dNTPs, 1 μL RNase Block). Samples were heated to 65 °C for 5 minutes, and incubated on ice for at least 2 minutes. 13 μL of each sample was removed to a new well as a control. 35 μL of the RT+ master mix (20 μL 5X 1st Strand Buffer, 5 μL 0.1 M DTT, 5 μL RNAse Block, 5 μL SuperScript III 200 U/μL) was added to each 65 μL sample. 7 μL of the RT-master mix (4 μL 5X 1st strand buffer, 1 μL 0.1 M DDT, 1 μL RNase Block, 1 μL water) was added to the remaining 13 μL of each sample. Samples were incubated at 50 °C for 2 hours, then were inactivated by heating at 70 °C for 15 minutes. Samples were stored at -20 °C.

To digest the RNA, first 1 μL DNase-free recombinant RNAse was added, and samples were incubated at 37 °C for 1 hour. For the RT-samples, we added RNAse diluted 1:5 in water, and added 1 μL to each well. A 1.5x SPRI bead cleanup was performed, where 150 μL of beads were added to the RT+ wells and 30 μL of beads were added to the RT-wells. Samples were washed 2x with 80% EtOH. Samples were eluted with water (145 μL for the RT+ well, and 22 μL in the RT-well), and 20 μL of each sample to a new well for PCR enrichment.

PCR reactions were set up on ice. 25 μL of PCR master mix (85 μL 5x GC Buffer, 42.5 μL 10 μL i7PCR primer, 12.75 μL 10 mM dNTPs, 8.5 μL 1 U/μL Kapa HiFi HotStart Polymerase, and 63.75 μL water) was added to each sample. 5 μL of 10 μM i5 index primer to each well in a column with a different primer for each column. cDNAs were enriched using the following PCR cycles: first 45 seconds at 98 °C, then 5 cycles of 15 seconds at 98 °C, 30 seconds at 64.5 °C, and 1 minute 10 seconds at 72 °C, then samples were held at 4 °C. qPCR survey reactions were set up using 5 μL of pre-enriched sample as a template to determine the optimal number of PCR cycles needed for each sample. The pre-enriched plates were kept covered on ice while the survey reaction ran. 9 μL of a master mix (2 μL 5X GC Buffer, 1 μL 10 μM i7PCR primer, 0.3 μL 10 mM dNTPs, 0.1 μL 1 U/μL Kapa HiFi HotStart Polymerase, 0.09 μL 100x Sybr, and 5.41 μL water) with an additional 1 μL of the i5 index to each well, for a final volume of 15 μL. The qPCR cycling conditions were as follows: 45 seconds at 98 °C, followed by 45 cycles of 15 seconds at 98 °C, 30 seconds at 64.5 °C, and 1 minute 10 seconds at 72 °C. Signal (R) was plotted against cycle number, and the number of cycles was determined by the necessary to reach ⅓ of the maximum R. The full samples were amplified by PCR using the following conditions: starting at 98 °C for 45 seconds, then the calculated number of cycles with 98 °C for 15 seconds, 64.5 °C for 30 seconds, and 72 °C for 1 minute and 10 seconds, and PCR was finished with 1 minute at 72 °C, and samples were held at 4 °C. A 9x SPRI bead cleanup was performed. The seven RT+ wells were eluted with 10 μL EB. 8 μL was collected per well, pooling elutions from the same column. The RT-wells were eluted with 14 μL EB. This solution was kept in the well with the beads bound to the magnet. QC was performed on these samples.

### Construction of pool–specific reference genome sequences

Aligning to a personalized genome can better represent alleles by avoiding biases in alignment to the reference genome^17,18,25^. Those previous methods were designed for sequence reads <100 bp. Paired end 250 bp reads cover substantially more variant positions in the human genome. For that reason, a novel custom alignment approach for longer paired end sequencing reads was developed.

Overall, our approach was based on the methods of Reddy et al, 2012 in that a personalized reference genome was created for short read alignment that consisted of a single sequence for each genomic region that lacks genetic variation, and two sequences for each genomic region that has genetic variants. Consecutive sequences were overlapped such that read alignments did not span multiple sequences. Here, that approach was extended to accommodate longer paired-end reads, and sequencing from a pool of five genomes.

First, a list of variants, including single nucleotide variants, insertions, and deletions, was obtained for each of the five individuals in our study using the Phase 3 release of the 1000 Genomes Project^29^. There were 11,291,340 single nucleotide variants, and 2,617,578 insertions or deletions. Next, phased sequences were created for all constant and variable regions in the pool of five individuals. Genetically constant regions were defined as those for which there is no heterozygosity in any individual in the pool; and genetically variable regions were defined as those for which there is at least one heterozygous genetic variant across the five individual genomes studies.

For each constant region, a single sequence of nucleotides was created with the sequence of that region, and all homozygous alternate variants were changed to match the genome sequence in our study population. For each variable region, a distinct sequence was created for each haplotype present in the study population. The reference sequence was modified in each haplotype to include the alternate alleles present in that haplotype. Haplotypes were inferred based on the 1000 Genomes Project phasing^29^. If the same haplotype occurred in multiple individuals in the pool, only one haplotype would be included in the custom reference genome.

To ensure all paired end reads from our STARR-seq libraries align to a single sequence on our reference genome, all constant regions were required to be at least 2 kb, and all variable regions were required to be at least 4 kb. Consecutive sequences were overlapped by 2 kb. If a variable region had more than 150 heterozygous or homozygous alternate variants that spanned more than 4 kb, the region was divided into two consecutive variable regions that overlapped by 2 kb.

After creation of the personal genome for the pool of individuals, the genome contained 43,734 constant and 135,000 variable regions. Each variable region contained on average 8.2 different haplotypes. The majority of the variable regions contained up to 112 different variants, with outliers in the variant count up to 291 variants. Each individual sequence was generally less than 100kb in length, with the longest sequences reaching almost 60Mb.

All code for creating personalized genomes is available at https://github.com/ReddyLab/Personal_Genomes.

### Alignment to pool-specific reference genome sequences

Bowtie 2 version 2.2.4 was used to align reads to the personal genome sequences^30^. First, a FASTA file was created with all personal genome sequences as described above. For each sequence in the FASTA file, the sequence name encoded the position and genotype of all genetic variants in that sequence. The genome was then indexed using bowtie2-build with default settings. The paired end reads were aligned to that indexed genome requiring no mismatches in the alignment seed, and paired ends were required to align to the same sequence in the expected orientation. The maximum fragment length was limited to 2 kb. Specifically, the options “bowtie2 -N 0 --no-mixed --no-discordant --dovetail -X 2000” were used. Finally, reads that occurred multiple times in the output libraries were removed based on unique molecular identifiers embedded in the sequencing reads using samtools rmdup command^31^.

### Identifying genomic regions with regulatory activity

Peak calling was performed using MACS on each of the three replicates individually, and resulting in three peak sets^32^. IDR correction was performed on each pair of peak sets. A single STARR-seq peak set was created from the union of these three corrected peak sets.

A set of ATAC-seq peaks was created using the union set of multiple experiments obtained from the ENCODE portal^16^.

Effect sizes of these peak sets were estimated using DESeq^33^. DESeq was used on the counts of reads in the ATAC-seq peaks directly. Size factors calculated from the STARR-seq peak set were not representative of the background genomic coverage in these datasets, and did not match the null hypothesis assumed by DESeq, resulting in calculated effect sizes that did not make sense in context. To remedy this issue, DESeq was used on the regions in the STARR-seq peak set from MACS^32^, with the size factors set to match those calculated with the ATAC-seq peak set.

### Characteristics of genetic variants that impact regulatory element activity

All regions containing variants identified in the first pool of samples were examined with the motifbreakR R package^21^. This provided both the position within the motif that included the location of this transcription factor, and estimated changes to transcription factor binding that would be caused by the specified variant.

These results were combined across similar transcription factor motifs which have been previously clustered by Vierstra^22^. When down sampling the individually identified transcription factor motifs to have only one occurrence per cluster, we chose the motif that had the greatest position weight matrix score. We focused our analysis on only concordant transcription factors, requiring that a transcription factor have a concordance between the estimated change in binding and the STARR-seq effect size at least 0.8, and at least 20 identified variants across the genome within that transcription factor (Supplementary Fig. 6). There were no transcription factor clusters with many identified variants and a concordance value below 0.2.

Information content analysis was performed on the individual transcription factors instead of the clustered transcription factors. The information content at each position was calculated using the following formula:

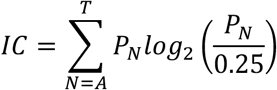

where IC is the information content value for that position, N is the nucleotide, and P_N_ is the probability of that nucleotide at that position. The percentage of variants in a single transcription factor motif were calculated, and compared to the information content at that position. The positions were also annotated as either having associated significant STARR-seq activity or having no associated STARR-seq activity. The information content in those categories was compared.

### Rare variants have greater impact on regulatory activity than common variants

Allele frequencies came from the gnomAD database^34^ for the African population. Trends in effect size versus population frequency were compared in two ways. First, a rolling average of the population frequency values and the effect sizes of the variants was plotted. Second, a subset of the variants based on their study frequency was taken, and linear regression comparing the population frequency and the effect size was performed. Finally, the variants were divided by four population frequency categories: common variants (AF>0.05), uncommon variants (0.01<AF<0.05), rare variants (0.001<AF<0.01), and ultra-rare variants (AF<0.01). The distributions of these categories were compared using a t-test.

### Multi-nucleotide insertions or deletions have greater impacts on regulatory activity than single nucleotide changes

Lengths of insertions and deletions were determined by the length of the longer allele. To compare distributions of effect sizes between SNVs and indels, we divided the dataset by the variant type. We plotted the length of the indel relative to the effect size of the variant.

### Correlation with chromatin QTLs

Chromatin accessibility QTL (caQTL) data from the African Functional Genomics Resource (AFGR) was used^23^. This dataset includes eQTLs, caQTLs, and sQTLs taken from a collection of multiple populations across Africa. These QTLs are associated each with an open chromatin region, which were used to subset the STARR-seq variants, and resulted in groups of one or more STARR-seq variants associated with each QTL. 1KGP data was used to calculate population-specific LD statistics where the sign of the statistic indicated the phase of the variants^35^. Combined STARR-seq effect sizes were calculated with their phased LD statistic value. The product of the log-transformed effect size of the STARR-seq variants across these groups was summed to find an aggregated effect size for the local STARR-seq variants.

To assess closed-chromatin STARR-seq effects, open chromatin regions were expanded up to 100kb. STARR-seq variants outside the open chromatin were treated identically to those inside the open chromatin in the correlations analysis.

---

Supplemental Tables are available at https://github.com/katherinedura/MiniSTARR.

**Supplementary Figure 1.**
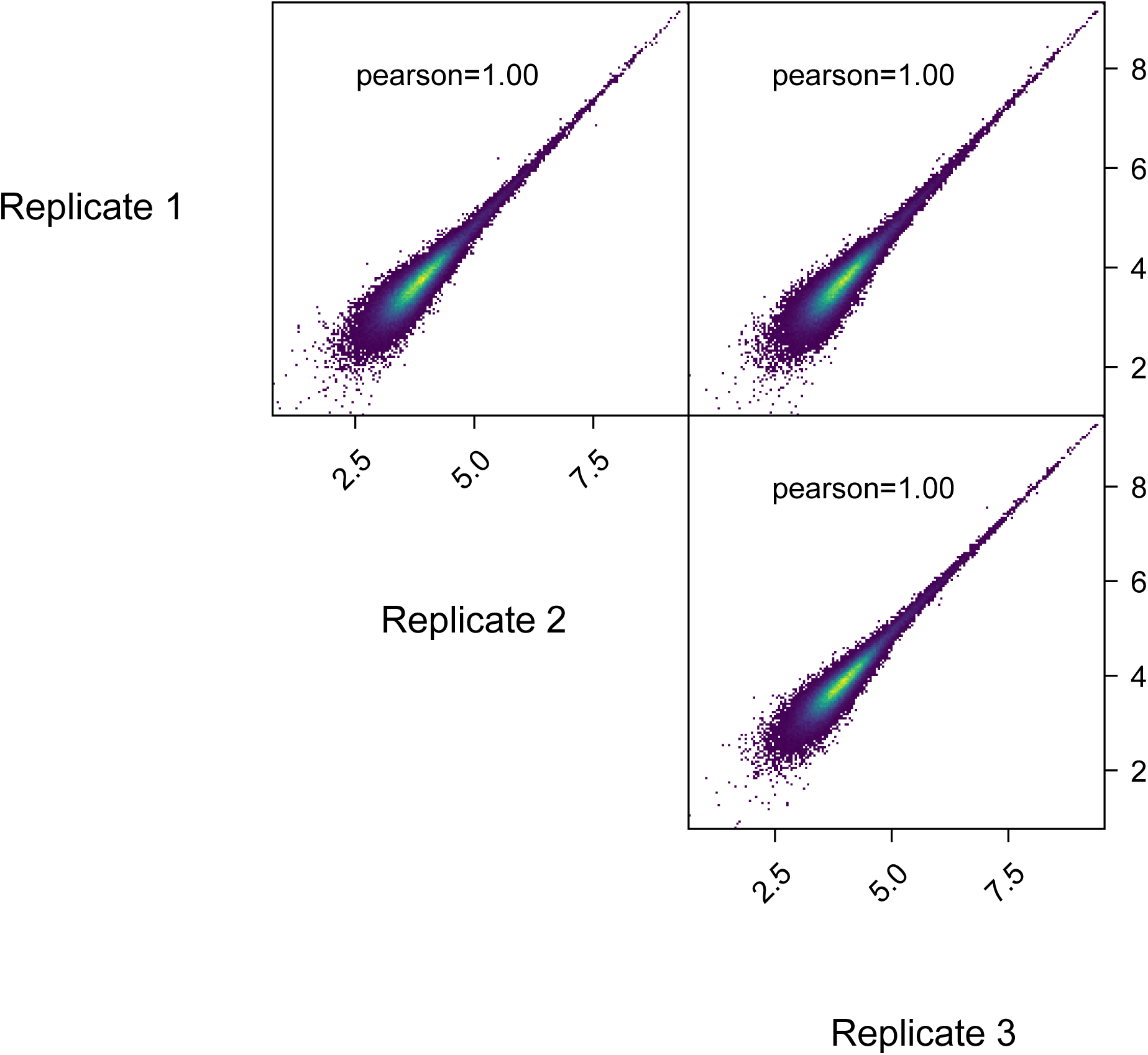
Correlation of coverage in the STARR-seq DNA libraries among three replicates.

**Supplementary Figure 2.**
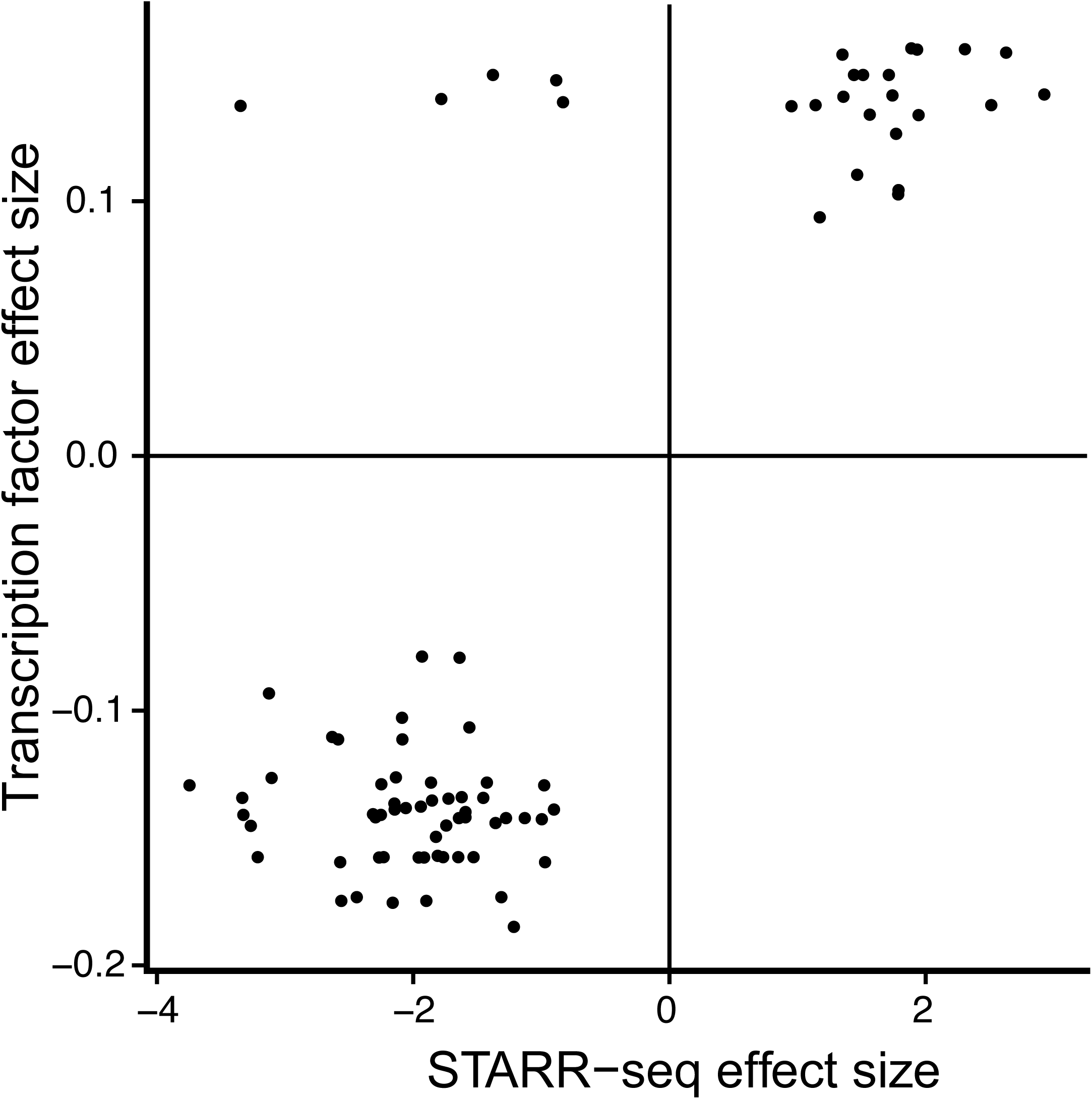
Correlation between STARR-seq effect size and change in AP-1 transcription factor binding.

**Supplementary Figure 3.**
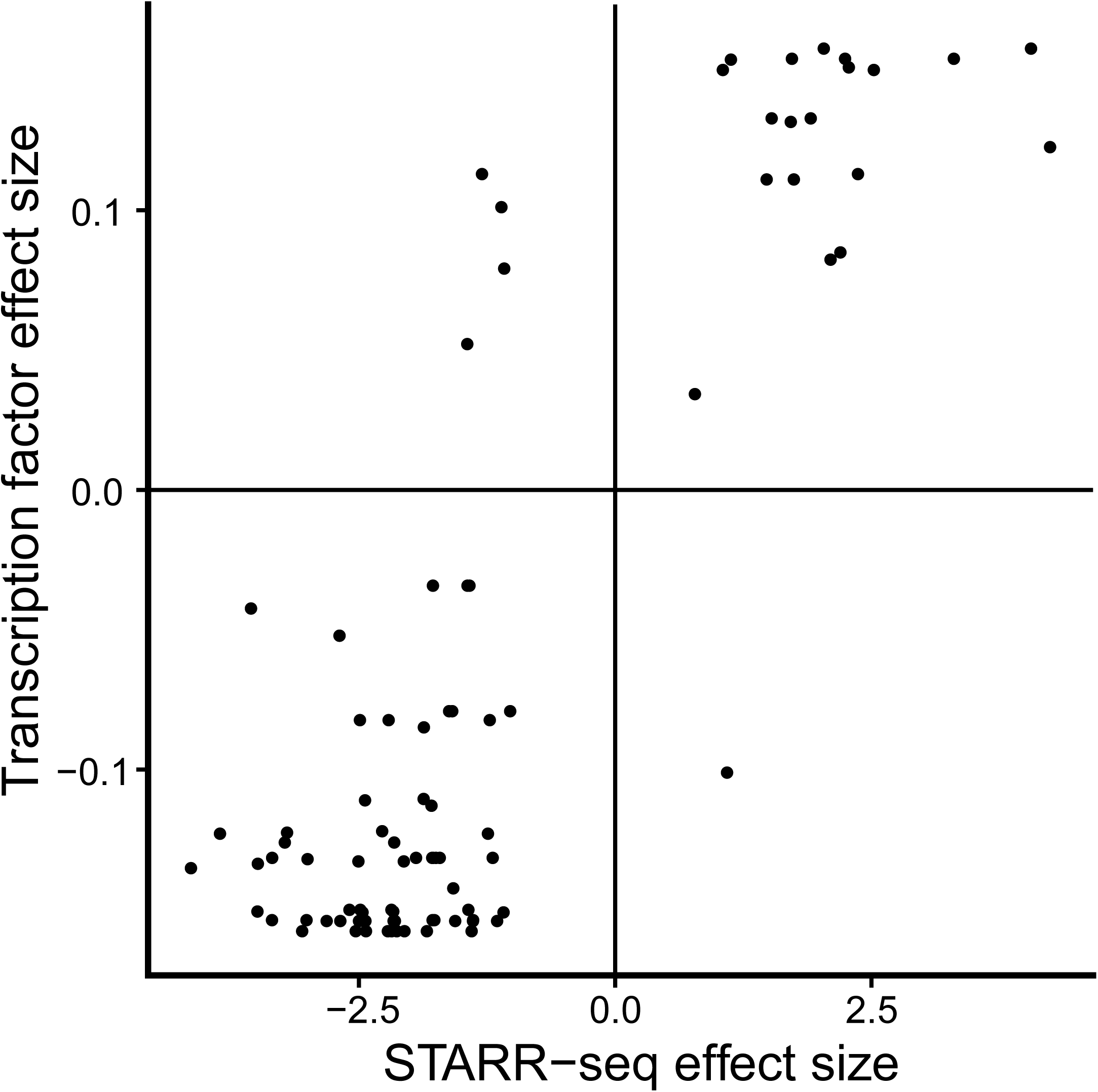
Correlation between STARR-seq effect size and change in CREB-AF3 transcription factor binding.

**Supplementary Figure 4.**
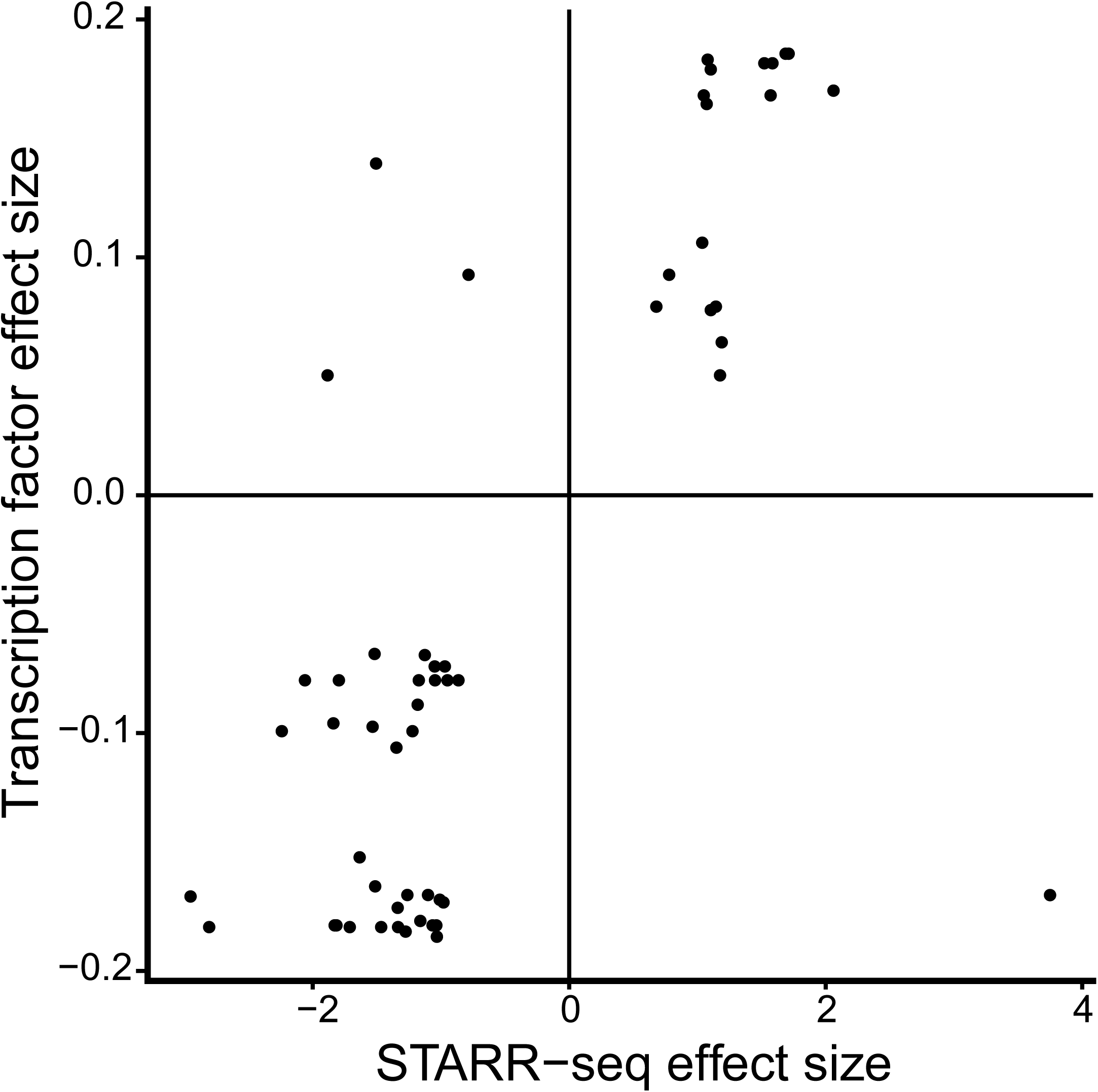
Correlation between STARR-seq effect size and change in ETS transcription factor binding.

**Supplementary Figure 5.**
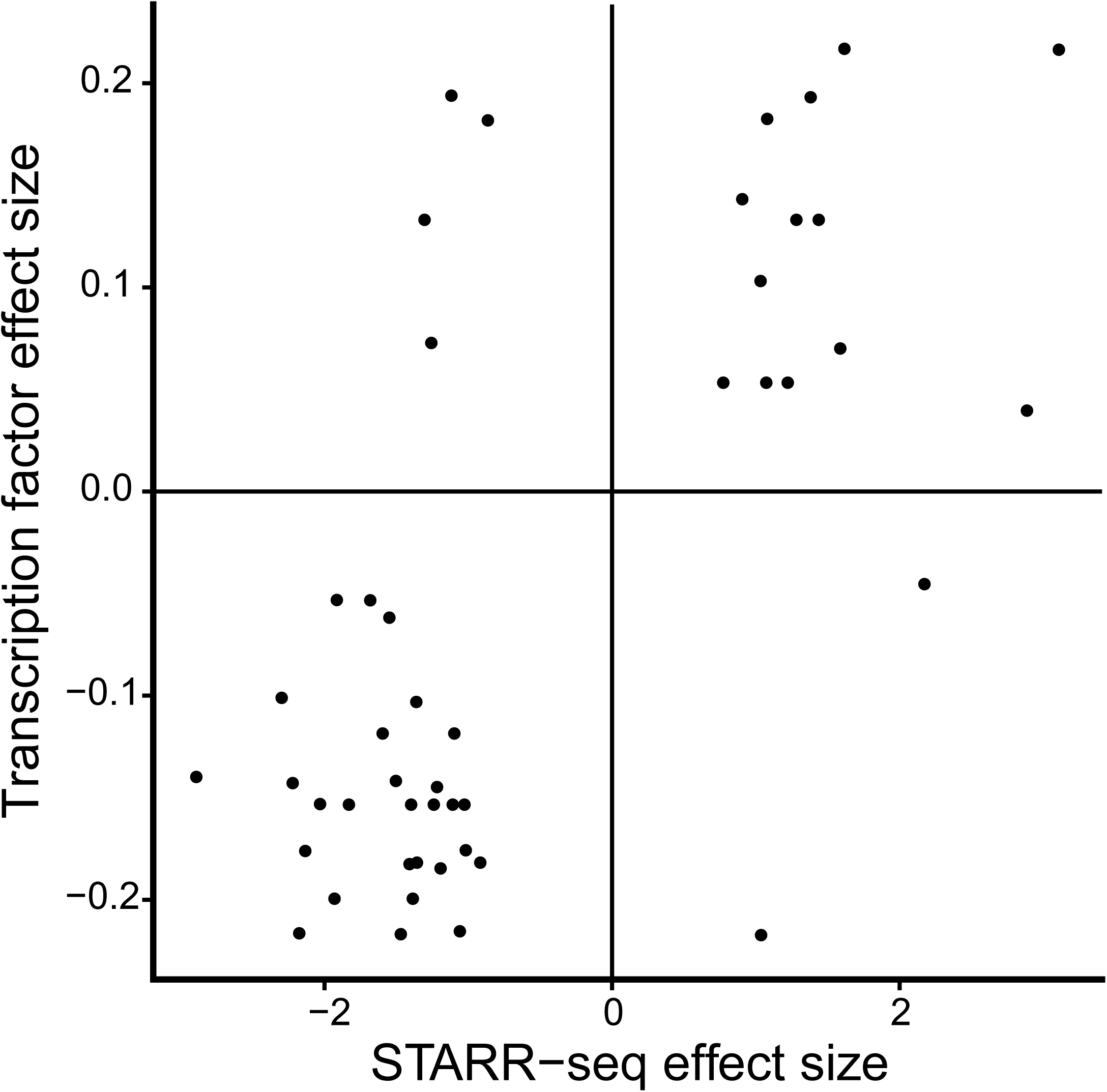
Correlation between STARR-seq effect size and change in GATA transcription factor binding.

**Supplementary Figure 6.**
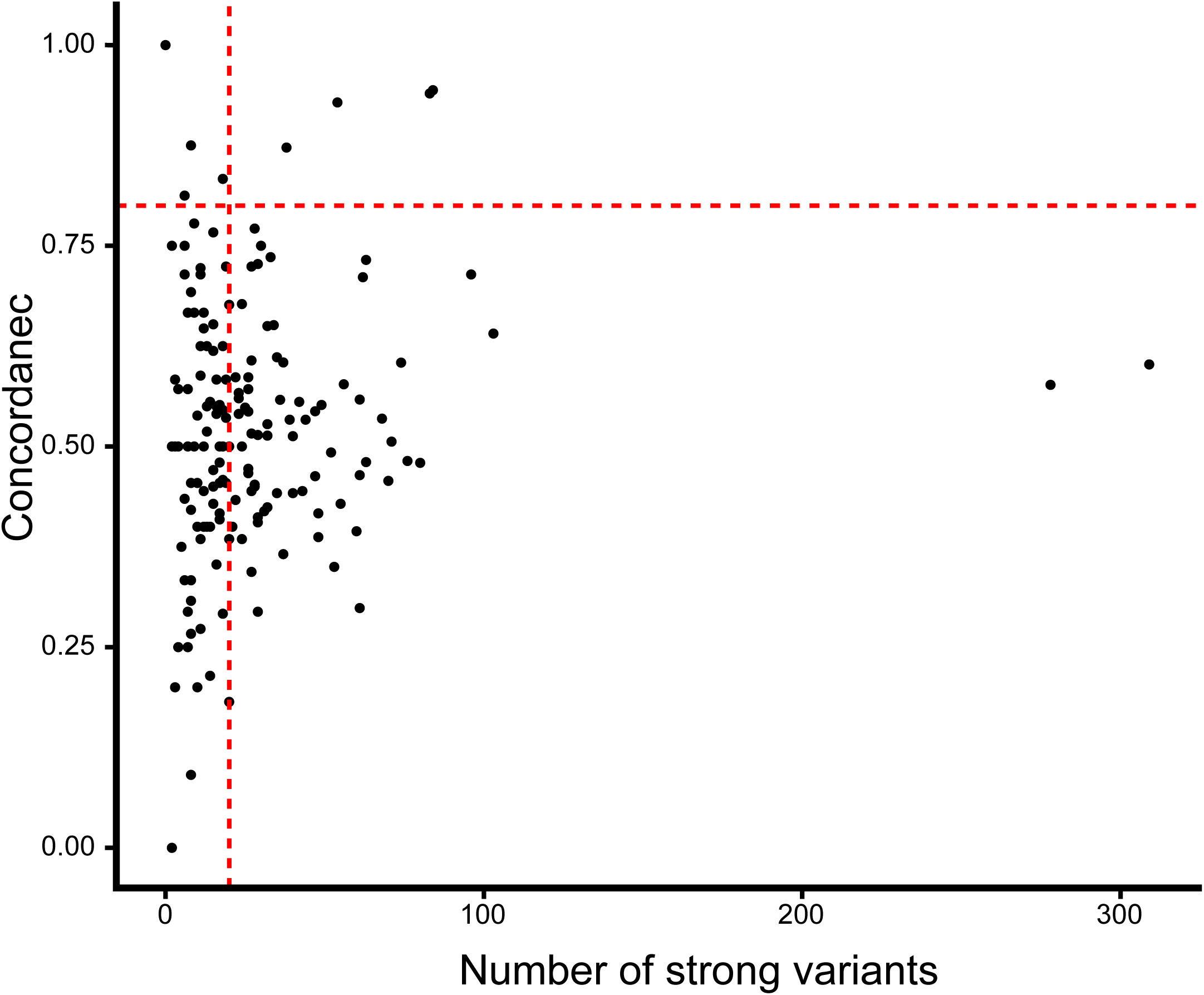
Selection criteria for inclusion of transcription factor motifs, including concordance between STARR-seq effect size and predicted change to binding above 0.8, and more than 20 identified variants with predicted strong effects.

